# Unique molecular characteristics of visceral afferents arising from different levels of the neuraxis: location of afferent somata predicts function and stimulus detection modalities

**DOI:** 10.1101/2020.06.06.138206

**Authors:** Kimberly A. Meerschaert, Peter C. Adelman, Robert L. Friedman, Kathryn M. Albers, H. R. Koerber, Brian M. Davis

## Abstract

Visceral organs receive neural innervation from sensory ganglia located adjacent to multiple levels of the brainstem and spinal cord. Here we examined whether molecular profiling could be used to identify functional clusters of colon afferents from thoracolumbar (TL), lumbosacral (LS), and nodose ganglia (NG) in the mouse. Profiling of TL and LS bladder afferents was also done. Visceral afferents were back-labeled using retrograde tracers injected into proximal and distal regions of colon or bladder, followed by single cell RT-qPCR and analysis via an automated hierarchical clustering method. Genes were chosen for assay (32 for bladder; 48 for colon) based on their established role in stimulus detection, regulation of sensitivity/function or neuroimmune interaction. A total of 132 colon afferents (from NG, TL and LS) and 128 bladder afferents (from TL and LS) were analyzed. Retrograde labeling from the colon showed NG and TL afferents innervate proximal and distal regions of the colon whereas 98% of LS afferents only project to distal regions. There were clusters of colon and bladder afferents, defined by mRNA profiling, that localized to either TL or LS ganglia. Mixed TL/LS clustering also was found. In addition, transcriptionally, NG colon afferents were almost completely segregated from colon DRG (TL or LS) neurons. These results indicate that populations of primary visceral afferents are functionally “tuned” to detect and interact with the internal environment and that information from all levels is integrated at higher (CNS) levels, not only for regulation of homeostatic functions, but for conscious visceral sensations including pain.

**Significance Statement:** Visceral organs are innervated by sensory neurons whose cell bodies are located in multiple ganglia associated with the brainstem and spinal cord. For the colon, this overlapping innervation is proposed to facilitate visceral sensation and homeostasis, where sensation and pain is mediated by spinal afferents and fear and anxiety (the affective aspects of visceral pain) are the domain of nodose afferents. Transcriptomic analysis performed here reveals that genes implicated in both homeostatic regulation and pain are found in afferents across all ganglia types, suggesting that conscious sensation and homeostatic regulation is the result of convergence, and not segregation, of sensory input.

## Introduction

Visceral organs receive sensory innervation from primary sensory neurons arising from multiple levels of the neuraxis. Depending on the organ, innervation can originate from vagal afferents (including afferents in the jugular and nodose (NG) ganglia), thoracolumbar (TL) spinal afferents (arising from lower thoracic and upper lumbar dorsal root ganglia (DRG)) or lumbosacral (LS) spinal afferents (arising from lower lumbar and upper sacral DRG). Hypotheses to account for why viscera requires monitoring by multiple populations of afferents include: a) that different levels of innervation are involved in different qualitative aspects of stimulus detection, including pain, b) that because the first synapse is on CNS neurons associated with sympathetic or parasympathetic circuits, different afferent populations provide the basis for integration of autonomic/homeostatic functions, or c) that the different levels play complementary roles in immune modulation.

With regard to shaping the nature of the visceral sensory experience, it had been hypothesized that spinal afferents, but not vagal afferents, relay nociceptive sensations (e.g., stabbing, burning, cramping), whereas vagal afferents underlie affective aspects of visceral pain including depression, anxiety, nausea and fear (Altschuler et al., 1993; Berthoud and Neuhuber, 2000; Grundy, 2002; Sengupta, 2009). However, recent evidence has shown that vagal afferents may have a significant role in nociception and pain (Lee et al., 2003; Kang et al., 2004; Yu et al., 2005; Bielefeldt et al., 2006; Canning et al., 2006; Nassenstein et al., 2008; Taylor-Clark et al., 2009). Even spinal afferents from different levels may have specific roles; colonic afferents arising from the TL level have been reported to be important for inflammatory pain but not acute pain, whereas LS colonic afferents are proposed to be involved in both acute and inflammatory pain (Traub et al., 1994; Traub, 2000; Wang et al., 2005).

In terms of homeostasis, spinal visceral afferents that innervate colon and bladder have cell bodies in both TL and LS DRG, projecting to organs they innervate along sympathetic (TL) or parasympathetic (LS) splanchnic nerves and terminate broadly in the spinal cord, including in the region of sympathetic (TL) or parasympathetic (LS) preganglionic neurons (Harrington et al., 2019; Smith-Edwards et al., 2019). In contrast, vagal afferents that innervate the colon (and possibly bladder (Herrity et al., 2014)) have cell bodies in the nodose ganglia, run in the vagus nerve with preganglionic parasympathetic fibers and terminate in the brainstem nucleus tractus solitarius (NTS) that in turn projects to nuclei in the brainstem, thalamus and cortex (Norgren, 1978; van der Kooy et al., 1984). Thus, redundant sensory input to the CNS could be informing different components of the autonomic nervous system.

Afferent/immune interactions are also critical for gut function and homeostasis, i.e., the vagus nerve has a crucial role in the anti-inflammatory reflex (Tracey, 2002), with vagal afferent response to cytokine release as the first step in this reflex. Primary afferents (vagal and spinal) release multiple peptides including calcitonin gene-related peptide (CGRP) and substance P that bind receptors on immune cells, allowing direct afferent and immune communication (Brain and Williams, 1985; Caceres et al., 2009; Altmayr et al., 2010; Riol-Blanco et al., 2014; Cohen et al., 2019). Ablation or silencing of primary afferents (vagal and spinal) also can modulate the immune response across multiple organs, including the lungs, small intestine and colon (Engel et al., 2011; Baral et al., 2018; Lai et al., 2020).

In this study we examined how genes required for sensation, homeostasis and neuroimmune regulation are divided among different visceral afferents. Single cell RT-qPCR analysis combined with automated hierarchical clustering (AHC) of afferents innervating the bladder and colon were carried out. Results show that location matters; that genes expressed by an afferent and where that afferent terminates (especially for the colon) is determined by the ganglionic location. Furthermore, the clustering of unique genes to afferents from different ganglia suggests that all afferents contribute to homeostatic and immune regulation, as well as signal all aspects of visceral sensation.

## Methods

### Animals

Experiments were conducted on male and female adult (6-12 week) C57BL/6 mice from Jackson Laboratory (JAX#000664). Animals were group-housed with a 12-hour light-dark cycle and *ad libitum* access to food and water. All procedures were approved by the Institutional Animal Care and Use Committee at the University of Pittsburgh and were carried out in accordance with AAALAC-approved practices.

### Back-labeling of neurons

Mice were anesthetized with isoflurane (2%) and a laparotomy was performed to access the pelvic viscera. Fluorescent retrograde dyes (cholera toxin subunit beta (CTB) or wheat germ agglutinin (WGA); Invitrogen) were injected into colon or bladder as previously described (Christianson et al., 2007). Briefly, using an insulin syringe (31-gauge needle), 5-10μL of Alexa Fluor 555-conjugated CTB was injected in 2-3μL aliquots beneath the serosal layer of the distal colon or at the base of the bladder. Alexa Fluor 488-conjugated CTB was injected in the proximal colon just distal to the cecum. To ensure dye did not spread between areas, colons were removed and visualized under a fluorescence microscope. In all cases, there was at least a 20mm area of middle colon that had no visible fluorescence. In a separate cohort of mice, 5-10μL of Alexa Fluor 488-conjugated CTB or WGA was injected in 2-3μL aliquots beneath the serosal layer in the urinary bladder body. WGA was used to examine whether CTB was labeling a specific subset of afferents (n=3, 3 female). No difference was detected in the afferents labeled with CTB or WGA therefore CTB was used in subsequent experiments. In a subset of these animals, 555-conjugated CTB was injected into the entire colon, combined with 488-CTB injection into the bladder to label convergent bladder/colon afferents. All incisions were then sutured, and animals were allowed to recover before returning to their home cage.

### Dissociation of ganglia and single neuron isolation

Three to five days after back-labeling, mice were anesthetized with isoflurane and transcardially perfused with cold Ca^2+^/Mg^2+^-free Hank’s balanced salt solution (HBSS). The NG, TL dorsal root ganglia (DRGs; T10-L1), and LS DRGs (L5-S1) were dissected into cold HBSS and enzymatically treated with cysteine, papain, collagenase type II, and dispase type II to facilitate isolation by mechanical trituration. Cells were plated on poly-d-lysine/laminin-coated coverslips in 35mmx10mm petri dishes. Coverslips were flooded with Dulbecco modified Eagle medium F12 (DMEM) containing 10% fetal bovine serum and 1% penicillin/streptomycin 2 hours later. Twenty-four cells were picked up from each culture using glass capillaries (World Precision Instruments) held by a 3-axis micromanipulator. Cells were transferred to tubes containing 3uL of lysis buffer (Epicentre, MessageBOOSTER kit), and stored at - 80°. Qualitative cell size measurement and relative number from different levels was noted, and effort was taken to sample cells of all sizes and level of innervation.

### Single cell amplification and qPCR

The methods for single cell RT-qPCR and clustering and validation of these techniques have been previously described (Adelman et al., 2019). Briefly, the RNA collected from each cell was reverse transcribed and amplified using T7 linear amplification (Epicentre, MessageBOOSTER kit for cell lysate), cleaned with RNA Cleaner & Concentrator-5 columns (Zymo Research), and assessed using qPCR with optimized primers and SsoAdvanced SYBR Green Master Mix (BIO-RAD). Cycle time (Ct) values were determined via regression. Quantification threshold for PCR was defined as the point at which there was a 95% replication rate (35 Ct) (Reiter et al., 2011). GAPDH threshold was thus defined as 25 Ct to ensure detection of transcripts at least a thousand-fold less common than GAPDH.

### Primer design and validation

Unique forward and reverse primer sequences were chosen for each gene within 500 bases of the 3’ end. Stock solutions of cDNA were generated by extracting RNA from the whole DRG and both 10 and 160pg aliquots of the RNA were amplified using the same procedure described above for single cells. Serial dilutions of these aliquots were used to calculate primer efficiencies over the range of RNA concentrations observed in single cells. Expression level was determined relative to GAPDH and corrected for these primer efficiencies (Pfaffl, 2001).

### Automated hierarchical clustering

All back-labeled neurons were clustered utilizing the unweighted pair group method with averaging (UPGMA) using expression information obtained from single cell PCR in MATLAB (MathWorks, Natick, MA). The preprocessing for this data analysis consists of taking the ΔCt values and replacing the samples that failed to generate a value for a given gene with the limit of detection for that gene.

### Tissue preparation and estimation of the number of back-labeled neurons

Three to five days after back-labeling, mice were isoflurane-anesthetized and transcardially perfused with cold isotonic saline. The NG, TL, and LS DRGs were removed and fixed with 4% paraformaldehyde solution for 30 minutes. For a negative control, the L3 DRG was also removed. After post-fixation, the ganglia were moved to 25% sucrose for overnight cryoprotection. After cryoprotection, ganglia were embedded in optimal cutting temperature compound (OCT; Fisher) and cryosectioned at 14 μm, allowed to dry, and immediately stored at −80°C. Each ganglion was placed on six slides with serial sections alternating between slides.

Ganglia sections were analyzed using fluorescent microscopy at 20×. Only two slides from the six were analyzed with 42μm between slides; visual confirmation was used to avoid analyzing the same cells more than once. At least 6 sections from each ganglion were analyzed per animal. For each animal, the number of 555-conjugated CTB positive, 488-conjugated CTB positive cells, and double labeled cells were counted. The percentage of CTB positive cells at each level of the neuraxis was found for each mouse and averaged within groups.

### Calcium imaging protocols

Colon neurons were back-labeled and prepared for culture as described above except proximal and distal colon were both labeled with CTB-555. Cells were incubated at 37°C and imaged within 36 hours as previously described (Malin et al., 2006). Prior to imaging, cells were incubated for 30 minutes at 37°C with 1 μl fura-2AM (2μM, Molecular Probes) and 2 μl of 20% Pluronic F-127 dissolved in DMSO (AnaSpec) and diluted in HBSS containing 10mg/ml BSA (Sigma). Coverslips were mounted on an inverted Olympus IX71 (Thornwood, NY) microscope stage with HBSS buffer flowing at 5ml/min, controlled by a fast step system (AutoMate Scientific). Perfusate temperature was maintained at 30°C using a heated stage and an in-line heating system (Warner Instruments). Chemicals were delivered with a rapid-switching local perfusion system. Firmly attached, CTB-positive neurons were identified using a 555nm filter and chosen as regions of interest using Simple HCI software (Compix Imaging Systems, Sewickley, PA). Unlabeled, adjacent cells were also identified and imaged. All fields were first tested with a brief application (4s) of 50mM K^+^ (high K^+^) to ensure that cells were responsive. Following a five-minute recovery period, agonists were applied in a randomized order with at least a five-minute recovery period between agonists. Stock solutions of capsaicin (Sigma) and mustard oil (Sigma) were made in 1-methyl-2-pyrrolidinone and α,β-methylene ATP (Sigma) was made in water. The stock solution of mouse interferon-α A (PBL Assay Science, Piscataway, NJ), recombinant mouse interleukin-4 (PeproTech, Rocky Hill, NJ), recombinant mouse interferon-γ and recombinant human interkeukin-8 (R&D Systems, Minneapolis, MN) was prepared in PBS containing 0.1% BSA, as previously described (Wang et al., 2017). Final concentrations applied to cells were 1μM capsaicin, 30μM α,β-methylene ATP, and 50μM mustard oil for four seconds and 500ng/ml for interleukin-4, interleukin-8 and interferon-γ and 1000iu/ml interferon-α for 90 seconds. Absorbance data at 340nm and 380nm were collected at one frame per second. Responses were measured as the ratio of 340/380nM excitation and 510nM emission over baseline (ΔF/F_0_; DG4, Sutter Instruments, Novato, CA). Peak responses were included in the analysis if the response was 5 standard deviations above baseline. The prevalence of responsive colon afferents was determined as a percentage of total healthy (high K^+^-responsive) CTB-positive cells. Any cell with significantly diminished Fura-2 signal over the duration of the experiment or that did not recover to baseline prior to the second agonist application was not included in the analysis.

### Experimental design and statistical analysis

For back-labeling experiments, the mean values of the number of back-labeled cells were compared between level (NG, TL, LS) and target (Proximal, Distal, Dual) using a two-way analysis of variance (ANOVA) with Tukey’s *post hoc* analysis. A two-tailed paired t-test was used to test for significance of the difference in the extent of spread (length) of CTB injections in the proximal and distal colon. For calcium imaging experiments, mean peak values of the calcium transients were compared between level (NG, TL, LS) and target (Proximal, Distal, Dual) using a mixed effects model with Tukey’s *post hoc* analysis. Statistical tests were performed in Excel (Microsoft, Redmond, WA) and GraphPad Prism (GraphPad Software, San Diego, CA). Data are expressed as mean ± standard error of the mean, where n represents mice used, unless indicated otherwise. The number of animals and statistical values are reported in the results section; significance is defined as *p* < 0.05.

## Results

### Distribution of back-labeled neurons from injection sites in distal and proximal colon

In mouse, the spinal innervation of the distal colon and bladder (including dually projecting colon and bladder neurons) have been described (Keast and De Groat, 1992; Wang et al., 1998; Robinson et al., 2004; Christianson et al., 2007), whereas innervation of the proximal colon has not been fully characterized. To fill this knowledge gap, the spinal and nodose sensory innervation of the proximal colon was investigated and compared to innervation of the distal colon.

Injection of CTB into the proximal and distal colon of individual mice labeled, on average, a total of 659.50 ± 75.36 CTB-positive neurons within the nodose and spinal ganglia (n=6, 4 males, 2 females; **Table 1**, **Fig. 1A**). There were nearly equal percentages of neurons arising from nodose (32.29% ± 3.73, *p* > 0.05), TL (30.29% ± 4.02) and LS (37.43% ± 5.77) ganglia. Injections into the distal colon produced more CTB-labeled cells (451.33 ± 83.78) compared to the proximal colon (181 ± 54.24, *F*_3,15_ = 22.19, *p* = 0.0256; **Table 1**), resulting in 65.42% ± 8.12 of CTB-positive afferents originating from distal colon, 27.87% ± 7.55 from proximal colon and 6.70% ± 1.02 being dual-labeled (i.e., innervating both proximal and distal colon). This difference in the number of cells innervating the proximal and distal colon occurs because the majority of LS afferents project to the distal colon; of the 258 + 56.23 back-labeled cells in LS ganglia, 251.67 + 58.70 were labeled by injections into the distal colon (this included 5.67 + 0.99 dual-labeled afferents; **Table 1**). Thus, LS colon afferents almost exclusively innervate the distal colon. For the proximal colon, the largest afferent contribution came from nodose and TL ganglia (58.29% ± 7.20 (NG), 38.22% ± 6.00 (TL), and 3.49% ± 1.63 (LS), *F*_4,20_ = 25.21, *p* <0.0001 for NG vs. LS, *p* = 0.0002 for TL vs. LS; **Fig. 1D**). The size of the proximal and distal injections was not a factor as the distance over which fluorescence could be detected in the proximal colon (13.0mm ± 1.9) and distal colon (14.0mm ± 1.1, *t* (7)=0.5739, *p* = 0.58) were equivalent (n=8, 5 males, 3 females; **Fig. 1E**). In addition, there was a 20-30mm gap between the leading edge of each injection. That TL afferents projected equally to proximal and distal colon, while LS afferents were more restricted to the distal colon is consistent with the idea that neural components that regulate colon function change along the length of the colon (Li et al., 2019).

**Table 1.**
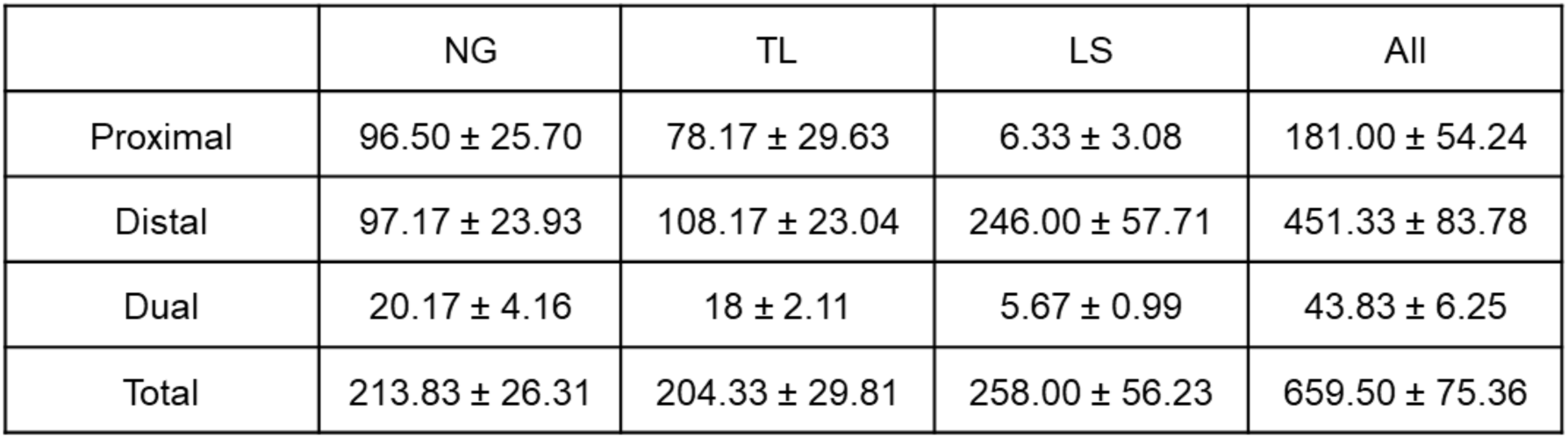
Number of NG (nodose), TL (thoracolumbar) and LS (lumbosacral) CTB-positive afferents projecting to proximal and distal colon. The total number of CTB back-labeled neurons from different levels of the neuraxis innervated proximal, distal or both proximal and distal colon (dual) almost equally. The NG and TL have approximately equal numbers of neurons that innervate proximal and distal colon, whereas LS cells almost exclusively innervate the distal colon.

**Fig. 1.**
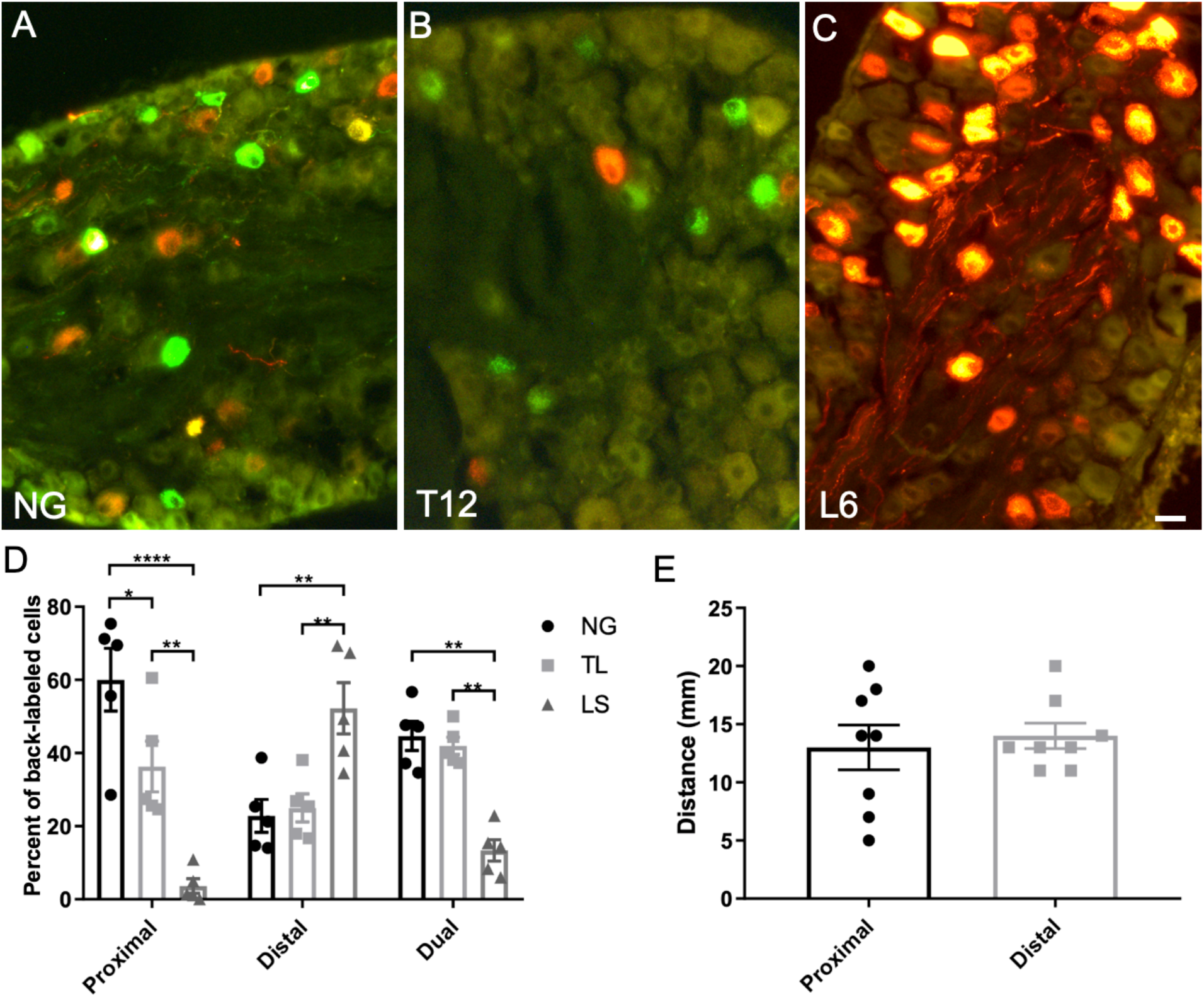
Proximal and distal colon have distinct patterns of innervation from different levels of the neuraxis. Retrograde labeled cell bodies in the NG (A), T12 DRG (B) and L6 DRG (C) following injection of CTB-488 (green) into the proximal colon and CTB-555 (red) into the distal colon. In all ganglia, the majority of cells contained one color of CTB. D) Quantification of the percentage of CTB positive cells across the neuraxis. NG and TL neurons project to proximal and distal colon, whereas LS neurons project primarily to distal colon. Afferents that project to both proximal and distal colon (dual) were most common in the NG and TL. E) Distance of fluorescence spread after CTB injection into the proximal and distal colon. The size of the injection ranged from 5 to 20 mm with 20-30mm between injection sites. Calibration bar A-C = 20μm. * *p*<.05, \*\**p*<.01, *** *p*=.001, \*\*\*\**p*<.001, using a two-way analysis of variance (ANOVA) (D) or paired two tailed t-test (E).

### Automated hierarchical clustering (AHC) based on mRNA expression and afferent location reveal distinct neuronal clusters

To classify afferent subtypes we used single-cell RT-qPCR to analyze transcripts of 28 genes previously used in a study for classification of identified cutaneous afferents (Adelman et al., 2019). These genes were shown to identify clusters that largely replicated the clusters produced by bulk, single cell RNAseq of DRG neurons unidentified with respect to innervation target or functional phenotype (Usoskin et al., 2015; Zeisel et al., 2018). Importantly, Adelman et al. combined AHC of these 28 genes with functional identification of neuron subtype with use of an *ex vivo* skin-nerve physiological preparation. This approach allowed correlation of molecular phenotype of individual cells with physiological response properties of intact afferents (e.g., conduction velocity, response to mechanical and thermal stimuli). Using this approach, these 28 genes produced clusters that corresponded to functional afferent subtypes (e.g., CMH (C-mechanoheat), CH (C-heat), and HTMR (high threshold mechanoreceptors)). For analysis of colon and bladder afferents, 4 additional genes encoding opioid receptors and the immune checkpoint inhibitor programmed cell death 1 ligand 1 (*Cd274*; AKA PDL-1) were added. For colon afferents only, 16 genes associated with immune function were also examined. Based on previous RNAseq analysis, these immune genes are highly expressed in both the nodose (Wang et al., 2017) and lumbar DRG (Usoskin et al., 2015).

The protocols used here were previously shown to produce a linear correlation between starting mRNA transcript level and the amount of mRNA detected following reverse transcription and amplification (Adelman et al., 2019). Thus, for each gene we can determine the relative level of expression for each cell analyzed (but cannot compare the level of expression between mRNA transcript species because of differential primer efficiency). Expression level information was converted into the heat maps shown in **Figs. 2 and 3**. In addition, descriptors of expression level in each cluster are reported as “undetected” (every cell in a group had no detectable level of transcript), “low” (when the average expression level for cells in a cluster was in the lowest 30^th^ percentile), “moderate” (between 31-70^th^ percentile) or “high” (71-100^th^ percentile).

**Fig. 2.**
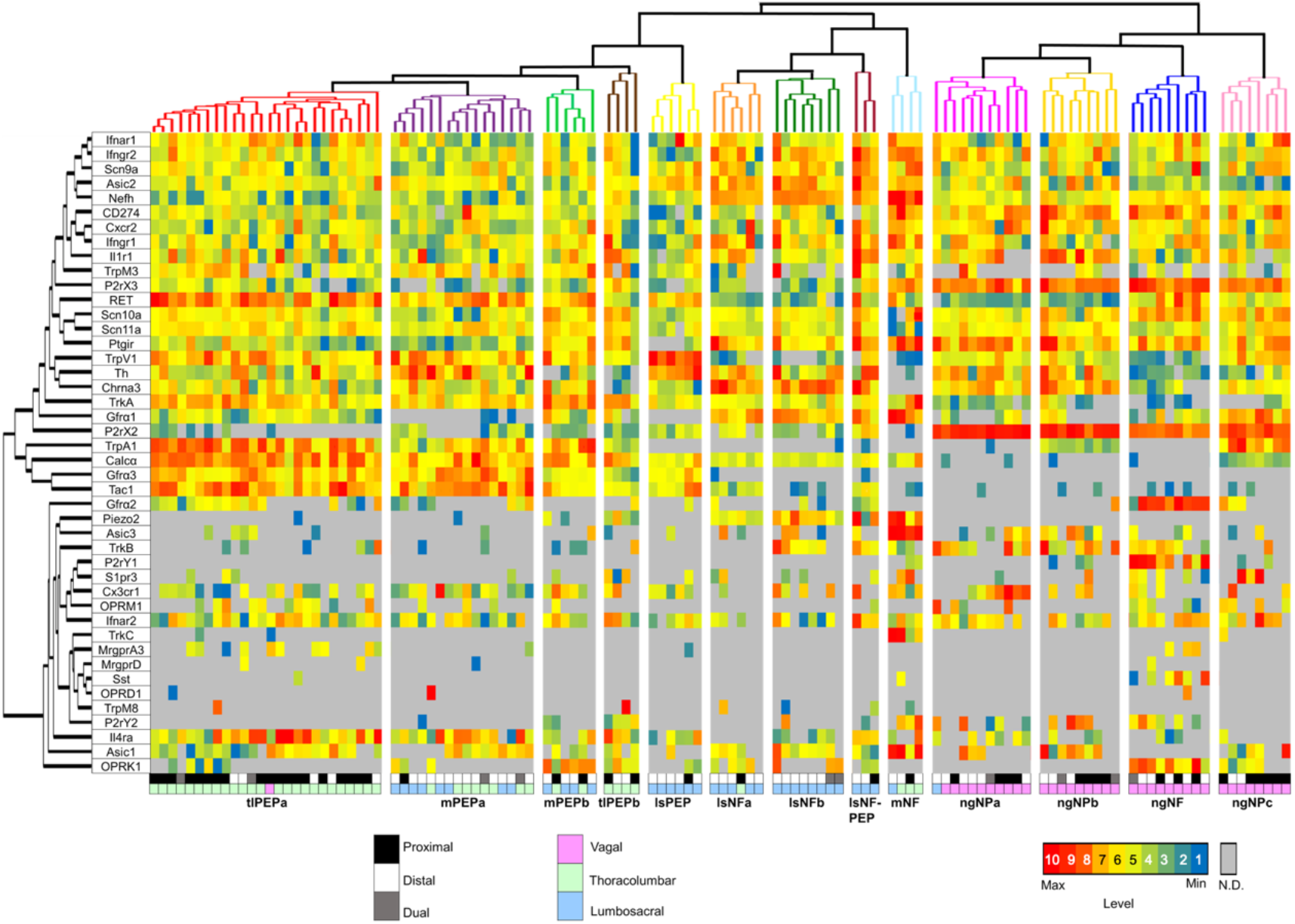
Colon afferents cluster into 13 distinct molecular clusters. Heatmap of mRNA expression across 116 colon afferents. Neurons and transcripts were clustered using the UPGMA Matlab algorithm. Red indicates high level of transcripts and blue the lowest. Gray bars indicate not detected (N.D.). The numbers in the color bar (bottom right) designate the colors that correspond to “low” (1-3), “moderate” (4-7) and “high” expression (7-10) used in the text. The bottom two rows of the heatmap indicate whether individual cells were back-labeled from injections into the proximal colon (black), distal colon (white) or proximal and distal (gray) and whether the cell body was in the NG (pink), TL (green) or LS (blue) ganglia. Cells grouped into clusters designated **tlPEPa** (# of cells=26), **mPEPa** (# of cells =16), **mPEPb** (# of cells =6), **tlPEPb** (# of cells 4), **lsPEP** (# of cells =6), **lsNFa** (# of cells =6), **lsNFb** (# of cells =8), **lsNF-PEP** (# of cells =3), **mNF** (# of cells =4), **ngNPa** (# of cells =11), **ngNPb** (# of cells =9), **ngNF** (# of cells =9), and **ngNPc** (# of cells =8).

**Fig. 3.**
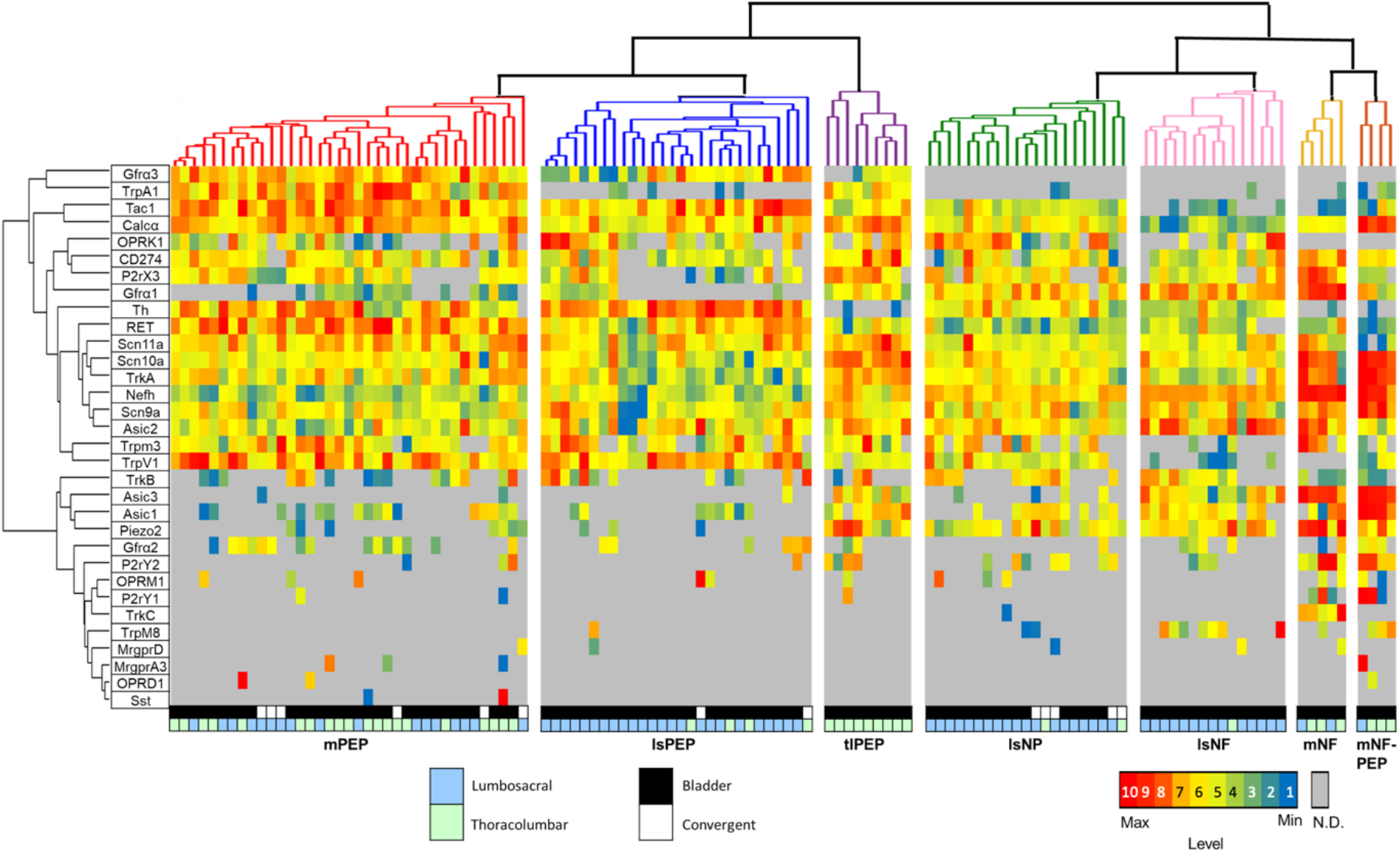
Bladder afferents cluster into 7 distinct molecular clusters. Heatmap of mRNA expression across 119 bladder afferents. Color coding for bladder afferents used the same methodology and parameters used for colon afferents. “Convergent” designates afferents that were back-labeled from retrograde makers placed in both colon and bladder. Bladder afferent groups were clustered into **mPEP** (# of cells 37), **lsPEP** (# of cells 28), **tlPEP** (# of cells 9), **lsNP** (# of cells 21), **lsNF** (# of cells 15), **mNF** (# of cells 5), and **mNF-PEP** (# of cells 4). As in the colon, some clusters arose from afferents located primarily at either TL or LS levels.

Clusters were named by combining the approaches of Zeisel et al. (Zeisel et al., 2018) and Hockley et al. (Hockley et al., 2018). Abbreviations used for clusters are “**PEP**” for peptidergic (clusters containing cells with expression of mRNA coding for substance P (*Tac1*) and calcitonin gene related peptide (*Calca*)), “**NP**” for nonpeptidergic (clusters of cells not expressing neuropeptide related mRNAs), “**NF**” for neurofilament (clusters of cells with expression of the high molecular weight neurofilament gene (*Nefh*), a marker of myelinated afferents (Lawson and Waddell, 1991; Adelman et al., 2019)), “**tl**” for clusters containing cells originating predominately (over 85%) from thoracolumbar ganglia, “**ls**” for clusters of cells from lumbosacral ganglia, “**ng**” for clusters of cells from nodose ganglia, and “**m**” for mixed clusters containing cells from TL and LS ganglia. Clusters are named based on whether they are **ng**, **ls**, **tl** or **m** plus whether they are **PEP**, **NP**, **NF** or some combination (e.g., **NF-PEP**). Because it is not possible to interrogate the unbiased algorithms used in this study, the contribution of each gene to the decision-making process is unknown. Therefore, genes that are highlighted in discussion of each cluster were chosen based on our inspection of the heatmaps and identification of ones most highly expressed in a cluster and ones that are either absent or expressed at low levels.

#### Colon afferents

Using the previously validated AHC methods of Adelman et al., (Adelman et al., 2019), a total of 132 colon afferents from NG (n=43), TL (n=48) and LS (n=41) separated into 13 distinct clusters based on mRNA level (n=7 animals, 4 males, 3 females; **Fig. 2**, **Fig. 4A**). 16 cells did not fit into any of these clusters and are not represented on the heatmap in **Fig. 2**.

**Fig. 4.**
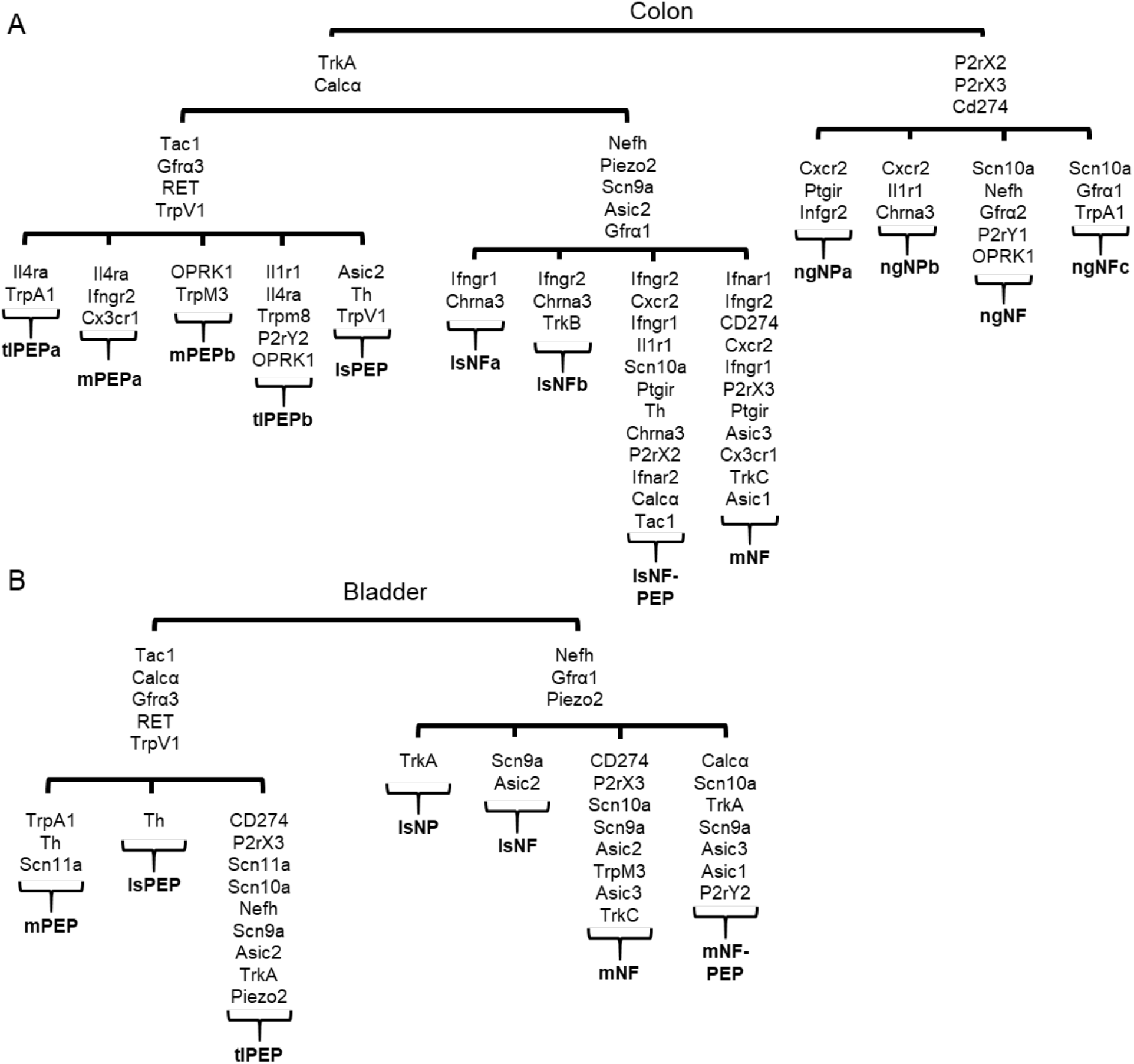
Colon and bladder afferent clusters have similar genes expressed but in different combinations. A) Schematic of major branch points from colon afferent clustering. Transcripts that differentiate each branch point are shown. B) Schematic of major branch points for bladder afferent clustering and the distinct genes in each cluster.

As the AHC is unbiased, it is interesting that the first branch point in the cell dendrogram separates nodose afferents from spinal afferents (**Fig. 2, Fig. 4A**). Major distinguishing characteristics of nodose afferents were the high levels of purinergic receptors (*P2rx2*, *P2rx3) and Cd274*, and low expression of calcitonin gene related polypeptide a (*Calca)* and preprotachykinin-1 (*Tac1)*. Nodose afferents were further divided into four clusters based on genes involved in growth factor receptor signaling, chemokine receptors and GPCR expression.

For AHC of spinal afferents, the first branch point separated cells based on levels of the heavy neurofilament (*Nefh*), a marker of myelination. The *Nefh* branch point gave rise to 5 *Nefh*-low and 4 *Nefh*-high final clusters. In the *Nefh*-low group, two clusters came almost exclusively from TL ganglia, one from LS ganglia, whereas two other clusters contained a mixture of TL and LS neurons. For the *Nefh*-high clusters, two contained only LS afferents and the other two contained a mix of TL and LS neurons. Thus, even at the initial level of analysis, the AHC revealed multiple mRNA-based clusters that were unique to afferents arising from anatomically distinct neuron populations.

### Nodose afferent clusters

Four nodose clusters with nearly equal number of cells (8-11 in each cluster) were identified (**Fig. 2, Fig. 4A**). Each cluster had both proximal colon and distal colon projecting cells. The **ngNPa** cluster has high levels of interleukin-8 receptor type 2 (*Cxcr2*), prostaglandin I2 receptor (*Ptgir*), and interferon gamma receptor 2 (*Ifngr2*). The **ngNPb** cluster also had high *Cxrc2*, interleukin-1 receptor type 1 (*Il1r1*) and cholinergic receptor nicotinic alpha 3 subunit (*Chrna3*). The **ngNF** cluster was a distinct nodose cluster with high expression of *Nefh*, glial cell line-derived neurotrophic factor family receptor alpha 2 (*Gfra2;* one of three co-receptors for *Re*t), sodium voltage-gated channel alpha subunit 10 (*Scn10a*), purinergic receptor *P2ry1* (a Gq coupled GPCR detected only with uniform, moderate expression in this one cluster), and opioid receptor kappa 1 (*Oprk1*). The nodose **ngNF** group was also unique in the uniform and low level of transient receptor potential cation channel family v member 1 (*Trpv1)* and *Chrna3*. The final nodose cluster (**ngNPc**) had high expression of *Scn10a* and glial cell line-derived neurotrophic factor family receptor alpha 1 (*Gfra1*, along with moderate expression of its co-receptor *Ret*) and was the only nodose cluster with a moderate level of transient receptor potential cation channel family a member 1 (*Trpa1*).

### Spinal Afferents - Thoracolumbar clusters

Of the two TL clusters (**tlPEPa** and **tlPEPb**), **tlPEPa**, overall the largest cluster, had high levels of *Calcα* and *Ret*, and moderate expression of glial cell line-derived neurotrophic factor family receptor alpha 3 (*Gfra3*; which was absent in nodose clusters), *Tac1*, tropomyosin receptor kinase a (*Trka*), *Trpa1*, *Trpv1* and interleukin-4 receptor alpha (*Il4ra*) (**Fig. 2, Fig. 4A**). The **tlPEPa** cluster neurons project primarily to the proximal colon. As anatomical origin (proximal vs. distal) was not a variable in the AHC algorithm, this clustering indicates that transcripts of proximal colon afferents are unique from those of the distal colon. The **tlPEPb** cluster, the second smallest of the spinal groups, had moderate *Il4ra*, low *Trpa1* and high *Trka*, relative to the **tlPEPa** cluster. This cluster also had high interleukin-1 receptor type 1 (*Il1r1*) and moderate transient receptor potential cation channel family m member 8 (*Trpm8*), purinergic receptor *P2ry2* and *Oprk1*.

### Spinal Afferents - Lumbosacral clusters

Within the four identified LS clusters, 20/23 of the back-labeled afferents were from the distal colon (two were dually labeled) indicating that the distal colon receives functionally distinct afferent input relative to the proximal colon. All but the **lsPEP** cluster had high or moderate *Nefh*, piezo type mechanosensitive ion channel component 2 (*Piezo2*), interferon gamma receptor 2 (*Ifngr2*) and *Gfra1*. The sodium voltage-gated channel alpha subunit 9 (*Scn9a*) was expressed in all LS neurons (including the **lsPEP** cluster) as was the acid sensing ion channel 2 (*Asic2*). The **lsNF-PEP** cluster was distinguished by high levels of the immune genes interferon gamma receptor 1 (*Ifngr1*), *Ifngr2*, *Cxcr2*, *Il1rl*, *Ptgir*, and interferon alpha receptor 2 (*Ifnar2*).

This cluster also had high *Chrna3* and moderate *Th*, *P2rx2*, *Tac1*, and *Calca*. The **lsNFa** and **lsNFb** clusters were the most similar of the LS clusters, distinguished by relatively high levels of *Ifngr1* in the **lsNFa** cluster and *Ifngr2* in the **lsNFb** cluster. The **lsNFb** cluster also had moderate tropomyosin receptor kinase b (*Trkb*), which was absent from other spinal clusters except the **lsNF-PEP** cluster. As mentioned above, the **lsPEP** cluster had transcripts not detected in other LS clusters and was unique in having the high levels of *Trpv1* and tyrosine hydroxylase (*Th*).

### Spinal afferents - Mixed clusters

There were 3 mixed clusters of spinal afferents that were defined by neurons within both TL and LS ganglia (**Fig. 2**, **Fig. 4A**). The **mPEPa** and **mPEPb** clusters had high or moderate *Calcα*, *Gfra3*, *Ret*, and *Trpv1* transcripts. The **mPEPa** cluster also had moderate levels of the immune genes *Il4ra*, *Ifngr2*, and fractalkine receptor (*Cx3cr1*). The **mPEPb** cluster was the only cluster with high transient receptor potential cation channel family m member 3 (*Trpm3*) and *Oprk1*, a gene with minimal expression in other spinal clusters (although present at high levels in one nodose cluster).

The other mixed cluster (**mNF**) was similar to the LS clusters with respect to the relative level of *Nefh*, Piezo2, Scn9a, *Gfra1*, and *Asic2*. This cluster also contained the immune related genes *Ifnar1*, *Ifngr2*, *Cd274*, *Cxcr2*, *Ifngr1*, *Ptgir*, and *Cx3cr1* as well as *P2rx3*, acid sensing ion channel 1 (*Asic1*), acid sensing ion channel 3 (*Asic3*), and tropomyosin receptor kinase c (*Trkc*). This was the only cluster (spinal or NG) that had no detectable level of *Chrna3*.

#### Bladder afferent clusters

For the bladder, 32 genes were analyzed from 128 back-labeled afferents from LS (80) and TL (48) levels (n=11 animals, 3 males, 8 females; **Fig. 3**, **Fig. 4B**). Neurons sorted into 7 distinct clusters with 9 cells not fitting into any cluster (not shown in the heat map). Because some afferents can project to both bladder and colon (Keast and De Groat, 1992; Christianson et al., 2007), in some experiments CTB conjugated to different fluorophores was injected into bladder and colon. From these animals (n=5, 2 males, 3 females) 14 convergent bladder/colon afferents were analyzed. Interestingly, these convergent afferents did not cluster into a separated group but were spread across the different clusters.

Unlike colon afferents, the first branchpoint in clustering of bladder afferents was defined by *Gfra3* and *Gfra1*. Neurons in these groups had either high levels of *Gfra1* or *Gfra3*, but never both. *Nefh*, the gene associated with the first branchpoint for colon afferents, was also highly segregated, being expressed at moderate to high levels in all *Gfra1* clusters, but moderately in only one *Gfra3* cluster. Similar to colon afferents, there were distinct TL and LS clusters as well as mixed clusters.

### Thoracolumbar cluster

Only one bladder afferent cluster was comprised exclusively of TL afferents (**tlPEP**) (**Fig. 3**, **Fig 4B**). The **tlPEP** cluster had high *Calcα*, *Cd274*, *Scn10a*, *Scn11a*, and *Trka* and moderate levels of *Nefh*, *P2rx3*, and *Piezo2*. This cluster was also distinctive in having both *Gfra1* and *Gfra3*, although at low to moderate levels. The low co-expression of *Gfra1* and *Gfra3* is likely the reason for positioning of this cluster at the border of the major branchpoint.

### Lumbosacral clusters

Bladder afferents formed three LS exclusive clusters. The **lsPEP** cluster had low *Gfra3* and almost no *GFRa1* (undetectable in 19/28 and moderate to low in the rest), high *Th* and moderate *Calca*, *Tac1*, Ret and *Tprv1*. The **lsNP** and **lsNF** clusters had no cells with detectable *Gfra3*, but moderate *Piezo2*. The **lsNP** and **lsNF** clusters can be separated by the high levels of *Asic2*, *Gfra1*, and *Nefh* in the **lsNF** cluster. The **lsNP** cluster also had high *Trka* (moderate in the **NP** vs. low in the **NF**).

### Mixed clusters

Bladder afferents formed three mixed clusters, **mPEP**, **mNF**, and **mNF-PEP**. The **mPEP** cluster was the largest and had the highest level of *Gfra3* of any bladder cluster. **mPEP** also had high *Tac1* and *Ret* and moderate *Calca, Trpa1, Trpv1, Th*, and *Scn11a*. The mixed **mNF** and **mNF-PEP** clusters were small and similar to the LS clusters based on *Gfra1* and *Nefh*, but also had high levels of several other genes including *Scn10a*, *Scn9a*, *Asic3*, and *Piezo2*. The **mNF** cluster can be differentiated from the **mNF-PEP** cluster by high *Cd274, P2rx3*, *Asic2*, *Trpm3*, and *Trkc*. In contrast, the **mNF-PEP** cluster had high *Calca*, *Trka*, *Asic1*, and *P2ry2*.

### Comparison of gene clustering for bladder and colon afferents

Comparison of high level transcripts across colon and bladder clusters shows many are present in afferents that innervate the two organs. However, different combinations of transcripts define the clusters (**Fig. 4A-B**). Dendrograms that illustrate the segregation of the 32 genes examined in spinal afferents for both organs (**Fig. 5A-B**) show some significant differences; the first major branch point for colon is associated with a peptidergic phenotype (e.g., *Calca*, *Tac1*, *Trka*, *Trpa1*, *Trpv1*, *Gfra3*) vs. a non-peptidergic phenotype (e.g., *Asic1, Trkb*, *Gfra2, Mrgprd, Mrgpra3, Trpm8, Piezo2*). The clustering in bladder shows a similar first branchpoint, however three high level and closely clustered genes *(Oprk1*, *Cd274* and *P2rx3)*, are shifted from the nonpeptidergic to the peptidergic group. In addition, in colon, *Trpv1* nearest neighbors are peptidergic-associated genes (*Gfra3* and *Trpa1*), whereas in the bladder *Trpv1* was separated from these genes by 3 branchpoints and clustered with *Trpm3*. Given that the overall percentage of cells expressing all 32 genes was not dramatically different for bladder and colon afferents, these differences suggest that at least some bladder afferents have novel properties not present in colon afferents.

**Fig. 5.**
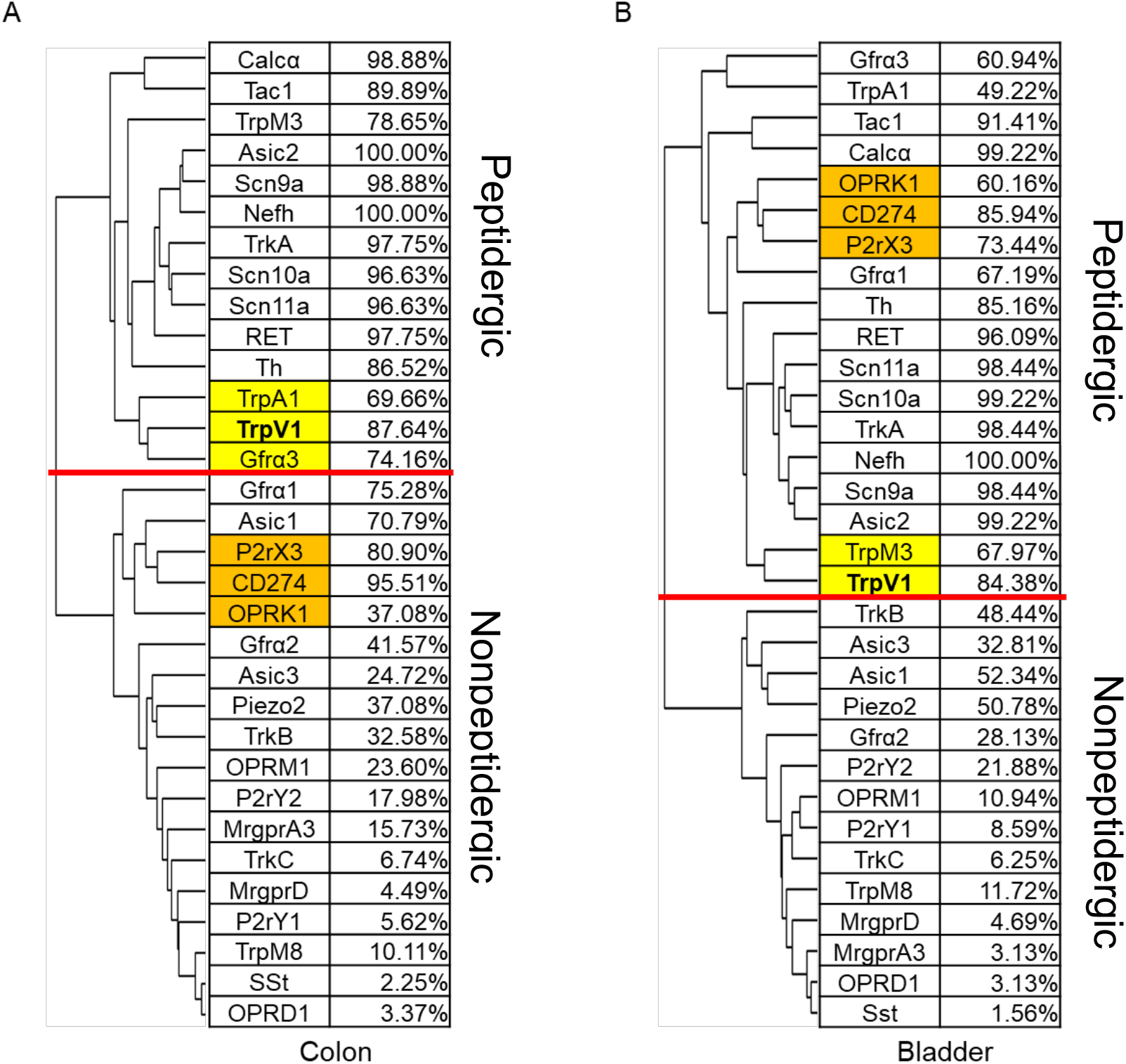
Spinal afferents of colon and bladder cluster into peptidergic and nonpeptidergic groups. A) Dendrogram of gene clustering for colon spinal afferents and percentage of cells with detectable transcripts. B) Dendrogram of gene clustering for bladder spinal afferents and the percentage of cells with detectable transcripts. Red line indicates the first major branch point in the dendrogram and separates peptidergic-from nonpeptidergic-related groups. Highlighted cells indicate differences between colon and bladder gene clustering.

### Calcium imaging and validation of gene function in colon afferents

Because RNA expression may not correlate with synthesis of functional protein (Adelman et al., 2019), we performed *in vitro* calcium imaging of back-labeled colon afferents to examine the relationship between mRNA expression and function. Afferents from different levels of the neuraxis were analyzed. Three agonists (capsaicin, mustard oil and α,β-methylene ATP) known to activate colon afferents (Christianson et al., 2010; Shinoda et al., 2010) via membrane bound receptors that were included in this analysis were used. We also examined the response properties of identified afferents to ligands of immune-related receptors (Interferon α, Interferon γ, Interleukin-4, and Interleukin-8) previously shown to produce calcium transients in primary afferents (Oetjen et al., 2017; Wang et al., 2017). The percent of responders was compared to the percent of cells that exhibited detectable (low, medium or high) levels of gene transcripts (**Table 2**).

**Table 2.** Calcium imaging of back-labeled colon afferents reveals relationship between mRNA level and receptor function. The percentage of cells that respond to selected agonists was compared to the percentage with detectable levels of the corresponding receptor gene(s). Capsaicin and mustard oil showed good agreement between responders and *Trpv1* or *Trpa1*, respectively. For α,β-methylene ATP, this was also true for NG afferents. TL and LS neurons exhibited many fewer responders, which is likely due to the reduced level of *P2rx2/3* in these cells. For immune-related receptors (*Ifnar1/2*, *Ifngr1/2, Cxcr2* and *Il4ra*), almost all afferents had detectable transcripts, but in most cases, fewer than half of the cells exhibited calcium transients in response to ligands. The number of cells analyzed is in parentheses.

|  | NG |  | TL |  | LS |  |
| --- | --- | --- | --- | --- | --- | --- |
| Agonist (Gene) | Percentage of Responsive Cells | Percentage of RNA expression | Percentage of Responsive Cells | Percentage of RNA expression | Percentage of Responsive Cells | Percentage of RNA expression |
| Capsaicin (*Trpv1*) | 74.28% (35) | 93.02% | 77.78% (18) | 95.83% | 95.45% (22) | 78.05% |
| Mustard Oil (*Trpa1*) | 36.58% (41) | 48.84% | 63.64% (22) | 91.67% | 47.37% (19) | 43.90% |
| $\alpha\beta$,methylene ATP (*P2rx2/P2rx3*) | 96.15% (52) | 97.67% / 97.67% | 34.78% (23) | 41.67% / 91.67% | 34.48% (29) | 75.61% / 68.29% |
| Interferon $\alpha$ (*Ifnar1/Ifnar2*) | 43.14% (51) | 95.35% / 76.74% | 18.18% (22) | 97.92% / 95.83% | 38.09% (21) | 100% / 80.49% |
| Interferon $\gamma$ (*Ifngr1/Ifngr2*) | 58.33% (48) | 95.35% / 97.67% | 34.78% (23) | 100% / 97.92% | 24.24% (33) | 100% / 100% |
| Interleukin-8 (*Cxcr2*) | 37.04% (27) | 97.67% | 15.38% (26) | 100% | 46.15% (26) | 100% |
| Interleukin-4 (*Il4ra*) | 41.18% (51) | 34.88% | 23.53% (17) | 91.67% | 47.83% (23) | 65.85% |

Calcium imaging was performed on 131 colon afferents (56 NG (n=6, 4 males, 2 females), 35 TL afferents (n=8, 2 males, 6 females), and 40 LS afferents (n=8, 4 males, 4 females); **Fig. 6A**). Many TL afferents were positive for *Trpa1* (91.67% vs. 48.84% for NG afferents and 43.90% for LS afferents) and this correlated with a high percentage of mustard oil responsive neurons (63.64%, compared to 36.58%, NG and 47.37%, LS). Capsaicin activated 74.28% of NG neurons, 77.78% of TL neurons, and 95.45% of LS neurons, consistent with the relatively broad *Trpv1* expression across all levels (**Table 2, Fig. 3**). Adelman et al. (Adelman et al., 2019) had shown for cutaneous afferents that neurons responsive to capsaicin had a ΔCt of 9 or lower for *Trpv1* (i.e., a ΔCt of 9 was the expression threshold). In this study, this level of *Trpv1* was met in approximately 80% of the colon afferents, roughly matching the percentage of neurons with capsaicin responses (**Table 2**).

**Fig. 6.**
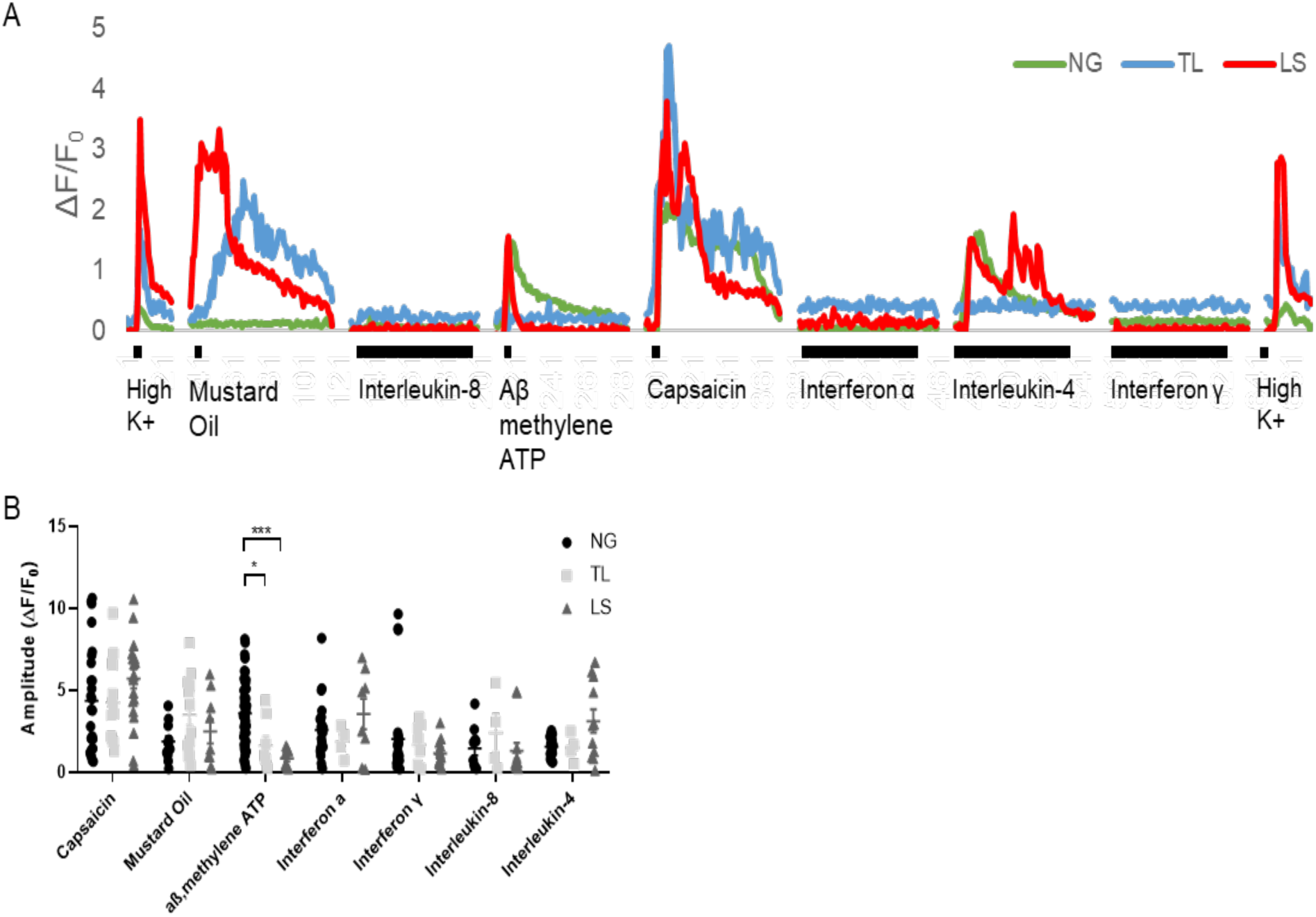
Colon afferents have functional receptors identified by single cell RT-qPCR. A) Sample trace of calcium transients from the NG, TL, and LS afferents. Black bar represents the length of agonist presentation and corresponds to 4 seconds (short bars) or 90 seconds (long bars). B) Amplitude of calcium transients from applied agonists from the different levels of the neuraxis. Amplitudes were similar for all agonists at all levels except for responses for α,β methyl ATP, which was significantly larger in NG afferents. \**p*<.05, \*\*\**p*<.001, using a mixed effects model.

The P2rX2/3 and P2rX3 selective agonist α,β-methylene ATP activated 96.15% of NG neurons, 34.78% of TL neurons, and 34.48% of LS neurons. For the NG, the percentage of α,β-methylene ATP responders (96.15%) matched the uniformly high level of *P2rx3* and *P2rx2* detected in NG afferents. In contrast, for TL and LS neurons, there were fewer responders to α,β-methylene ATP (34.78% and 34.48%, respectively) relative to the percentage of neurons expressing detectable levels of *P2rx3* and *P2rx2 (see **Table 2**)*. Adelman et al. found that for cutaneous afferents expressing the *P2rx3* receptor (*P2rx2* was not examined), a ΔCt of 6 was the threshold for presence of functional receptors. This value was measured in only 20.45% of TL neurons and 14.28% of LS neurons, consistent with the low percentage of afferents responding to α,β-methylene ATP. In terms of the amplitude of the calcium transients in response to α,β-methylene ATP, for neurons that did respond, the peak responses of NG afferents was significantly higher than for LS and TL afferents (*F*_12,210_= 3.065, *p* = 0.0204 for TL, *p* < 0.0001 for LS; **Fig. 6B**). This suggests that for P2X receptors, there is a direct relationship between the transcript level and response to ligand.

With the exception of *Il4ra* in NG and LS neurons, all immune-related receptors assayed were detected in nearly all cells. Unlike other ligands examined, the percentage of neurons activated by the immune-receptor ligands was significantly lower than the percentage of cells positive for these receptor genes (except for *l4ra* in the NG). In NG neurons, calcium responses to interferon-α, interferon-γ, interleukin-8, and interleukin-4 occurred in approximately half (37-58%) of the cells expressing the corresponding receptor mRNA. A lower percentage of TL and LS neurons exhibited calcium transients in response to these ligands (24-47%, LS and 15-34%, TL neurons). These results indicate that mRNA alone cannot be used as an indicator of functional activity for these receptors. For cells where immune ligands did cause a calcium transient, the amplitude of these transients were similar to those for capsaicin, mustard oil, and α,β-methylene ATP (**Fig. 6B**).

## Discussion

Identification of unique protein or mRNA markers that can be used as surrogates for functional phenotyping has long been a goal of primary afferent biology. Here we employed single cell mRNA analyses of up to 42 genes to determine if a unique molecular signature for colon and bladder primary afferents mapped to their origin, i.e, NG, TL or LS ganglia. Using unbiased, automated hierarchical gene clustering, we found that for both colon and bladder afferents, most clusters had neurons primarily from a single level of the neuraxis (>85% of cells from NG, TL or LS) than from mixed levels (**Fig. 7**). For the 13 colon clusters, 4 originated in the NG (**ngNPa-c, ngNF**), 2 from TL (**tlPEPa&b**) and 4 from LS (**lsPEP, lsNFa&b** and **lsNF-PEP**). Of the 7 bladder clusters, 1 came from TL (**tlPEP**) and 3 from LS (**lsPEP, lsNP**, and **lsNF**) ganglia (**Fig. 7**). For colon, there were also clusters that primarily innervated the proximal (**tlPEPa**) or distal (**mPEPa, lsPEP, lsNFa&b**) colon. Genes in our panel were chosen based on a validated role in stimulus detection (e.g., TRP channels, piezo2, P_2_X receptors, ASIC channels, cytokine receptors) or responsiveness (e.g., opioid receptors, neuropeptides, sodium channels), suggesting that function-specific components of sensory information arriving in the CNS is segregated by sensory ganglia at different levels.

**Fig. 7.**
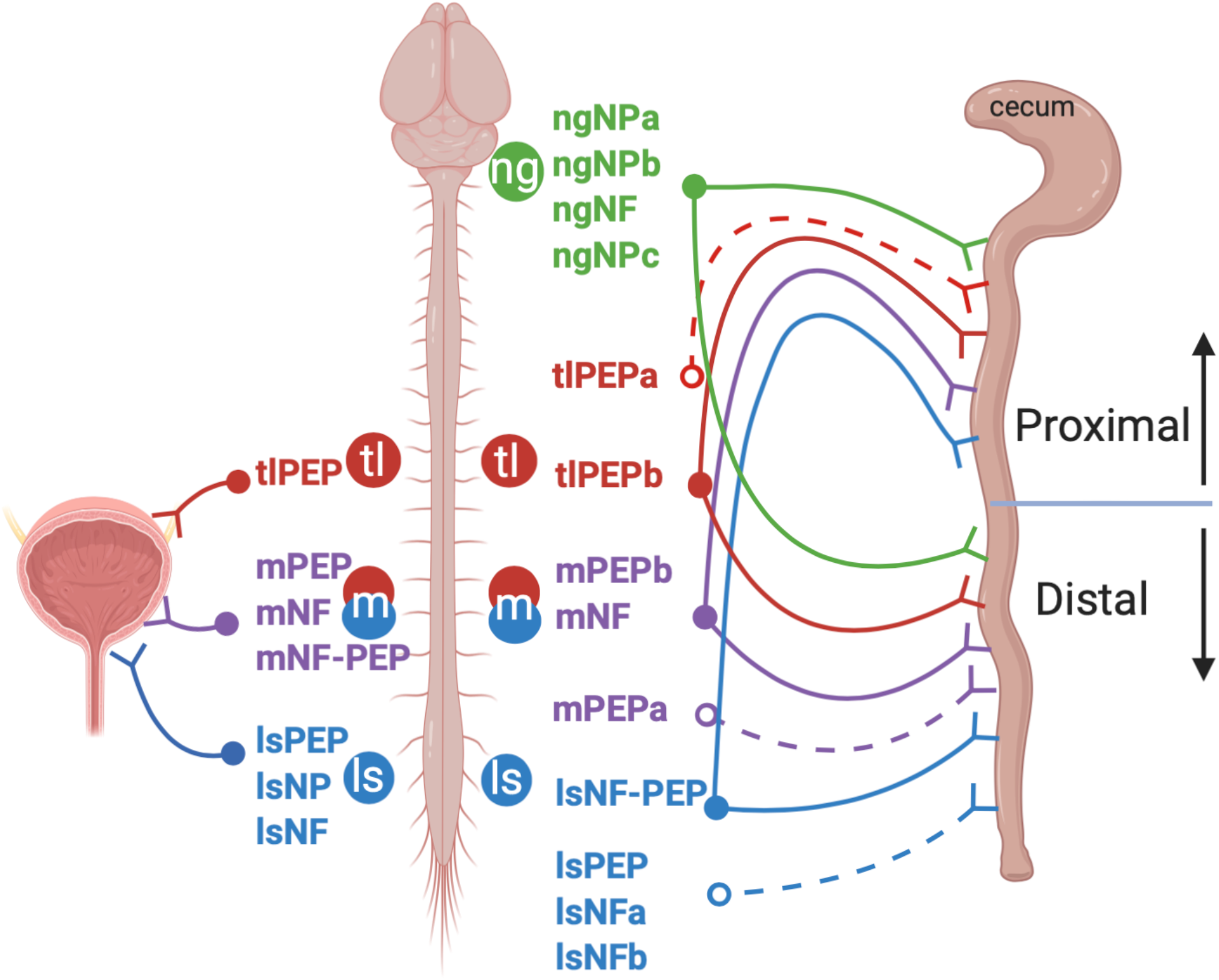
Automated hierarchical clustering shows afferent clusters for colon and bladder arise from distinct sensory ganglia. For both bladder and colon, the majority of the identified clusters are from one anatomical level; i.e., not from mixed (m) ganglia (11/13 for colon and 4/7 for the bladder). For the colon, there are also clusters that project selectively to either the proximal or distal colon (dotted lines). Illustration created with BioRender.com.

Unlike RNAseq analysis, our study focused on a small number of curated genes. The clusters identified are similar to those detected in a previous RNAseq analysis of back-labeled distal colon afferents from both TL and LS levels (Hockley et al., 2018). Using single-cell consensus clustering (Kiselev et al., 2017) of 314 cells, 7 clusters were identified based on the presence or absence of peptidergic (*Calcα, Tac1, Trpa1*) or nonpeptidergic (*Mrgprd, P2rx3, Gfrα2*)) markers, and the expression of *Nefh* (which was coexpressed with *Piezo2*). Hockley et al., also identified mixed clusters arising from a combination of TL and LS afferents and clusters that were exclusively from LS afferents (defined as “pelvic nerve” afferents).

A significant finding of the current study is the nearly exclusive manner in which colon NG neurons segregate from colon spinal afferents. Only one colon DRG neuron appeared in the 4 nodose clusters and only one nodose neuron sorted with neurons in the spinal clusters. The major contributor to this segregated pattern was the uniformly high levels of *P2rx2* and *P2rx3* in NG neurons. Similarly high levels of the prostacyclin receptor (*Ptgir*) and *Cd274* (AKA PDL-1 (programmed cell death ligand 1)) were in the NG. *Ptgir* and *Cd274* have important roles in immune responses, consistent with the role of nodose afferents in neuroimmune regulation (Sharkey and Mawe, 2002; Tracey, 2002; Baral et al., 2018).

The individual NG clusters were also unique. No NG cluster had characteristics similar to the peptidergic clusters at TL or LS levels. There was a conspicuous absence or low level of *Trka, Calca, Gfra3 and Tac1* in NG neurons, consistent with previous findings (Wang et al., 2017; Trancikova et al., 2018). The NG clusters were also unique in that unlike TL or LS clusters, NG clusters projected equally to proximal and distal colon with the overall number being virtually the same (45.13% proximal vs. 45.44% distal, compared to a 38.25%/52.94% (proximal/distal) split for TL ganglia and 2.54%/95.35% split for LS ganglia). This suggests that unlike clusters arising from DRG afferents, innervation by NG afferents of different clusters is required throughout the length of the colon. This could reflect a role for NG afferents in neuroimmune monitoring, which would be in line with the high percentage of NG neurons that exhibited calcium transients in response to cytokine application.

RNAseq studies have been performed on NG afferents and compared to jugular afferents (Wang et al., 2017; Kupari et al., 2019). In mouse, jugular afferent somata are located in a ganglion fused to the placode-derived NG and like DRG neurons, are neural crest derived (Baker and Schlosser, 2005)). Wang et al. and Kupari et al. found that like the DRG neurons examined here, jugular neurons are molecularly distinct from NG afferents (Wang et al., 2017; Kupari et al., 2019). However, cluster analysis found several more unique NG clusters then identified in the present study (18 nodose clusters with a major breakdown between “mechanosensitive” groups (1-11) and “nociceptive” clusters (12-18); (Kupari et al., 2019). A likely reason for this difference is that we used back-labeled neurons from a single organ (colon) whereas in Kupari et al., clusters from all the thoracic and abdominal organs innervated by the NG were represented.

A major theory to explain the need for overlapping sensory innervation of the viscera, especially for those organs that receive nodose and spinal afferent input like the colon, is that different pathways relay unique aspects of the sensory experience. For example, in the case of pain arising from abdominal organs (including the colon), spinal afferents have been proposed as the transducer of noxious pain (e.g., sharp, burning) whereas nodose afferents transmit affective aspects of visceral pain (e.g., fear, anxiety, nausea) (Berthoud and Neuhuber, 2000; Grundy, 2002; Sengupta, 2009). These distinctions are supported by patient reports of sensations produced by whole vagal nerve stimulation (Sackeim et al., 2001; Ben-Menachem, 2002), but contradicted by the similarities in firing properties of nodose and spinal afferents innervating the gut (Sengupta et al., 1990; Ozaki and Gebhart, 2001; Yu et al., 2005; Bielefeldt et al., 2006) and the anecdotal reports of noxious gut pain in patients with spinal transections, i.e., patients that lack ascending input from spinal afferents (Charney et al., 1975; Yung and Groah, 2001; Finnerup et al., 2008; Levinthal and Bielefeldt, 2012)).

Given that in most cases activation of visceral afferents never produce conscious sensations (Critchley and Harrison, 2013), the data presented here could be used to argue that the driving force behind the evolution of the complex pattern of sensory innervation arises out of the requirements of homeostatic regulation and not because the different afferent populations are “labeled-lines” carrying segregated qualities of visceral sensation. First, the projections of NG and TL clusters overlap anatomically in a manner that suggests these molecularly distinct afferent populations are making unique contributions to homeostatic function; both NG and TL afferents innervate the entire colon, with approximately equal numbers of afferents, providing direct input to their associated parasympathetic (NG) and sympathetic (TL) CNS circuits in the spinal cord and brainstem. Second, while it is clear that neurons at NG, TL and LS levels have unique transcriptomic profiles with respect to receptor expression (e.g., TRP and purinergic channels, cytokine receptors) that should “tune” them to different aspects of the internal milieu, all clusters, regardless of origin, express receptors required for monitoring of homeostasis-relevant stimuli. Third, many organs, in particular the small and large bowel, contain large numbers of innate immune cells and their interactions with their local microbiome affect the entire body (Cryan and O’mahony, 2011; Maynard et al., 2012). The near ubiquitous expression of immune-related genes in all clusters is consistent with the idea that these afferents play a role in integration of information required for immunological homeostasis. Moreover, the efferent function of visceral afferents, i.e., their ability to release molecules (e.g., peptides likes CGRP and substance P) that can modulate immune cells has recently been shown to play a central role in somatic and visceral tissue responses to pathogenic insult (Baral et al., 2018; Lai et al., 2020; Saloman et al., 2020).

The corollary to this hypothesis is that different aspects of conscious visceral sensation (e.g., both noxious and affective aspects of visceral pain) is the result of integration of information gathered from all sensory ganglia and then processed by different (but likely overlapping) CNS circuits. Five colon afferent fiber types have been identified: mucosal, muscular, serosal, mesenteric, and muscular/mucosal (Brierley et al., 2004; Brierley et al., 2005). Both TL and LS have mucosal, muscular, and serosal afferents whereas mesenteric afferents are specific to TL neurons and muscular/mucosal afferents are specific to LS afferents. This suggests that many TL and LS afferents detect similar mechanical stimuli, whereas others code for unique aspects of these stimuli. That neurons from all levels of the neuraxis (including the NG) have a role in conscious sensation including pain is supported by the data presented here; all of the colon clusters express proteins shown to play important roles and/or are “required” for nociception in both somatic and visceral afferents (Delafoy et al., 2006; Bielefeldt and Davis, 2008; McIlwrath et al., 2009; Kiyatkin et al., 2013). It seems unlikely that noxious pain would be detected by only a subset of these afferents from one or two levels with the remainder transmitting signals specific to non-noxious qualities of pain (i.e., affective aspects). The more parsimonious explanation is that afferents in all clusters, at all levels, are collaborating to provide a comprehensive report of peripheral conditions and occasionally this information reaches qualitative and quantitative thresholds required for conscious visceral sensation, including pain.

All authors have no conflict of interest.

## Acknowledgements

The authors acknowledge and thank Mr. Christopher Sullivan for expert technical support and mouse husbandry.

## Author contributions

KAM, conception and design, acquisition of data, analysis and interpretation of data, drafting and revising the article. PCA and RLF, acquisition of data, analysis and interpretation of data. KMA, HRK and BMD, conception and design, analysis and interpretation of data, drafting and revising the article.

## Abbreviations

DRG: dorsal root ganglia
NG: nodose ganglia
RT-qPCR: real-time quantitative polymerase chain reaction
TL: thoracolumbar
LS: lumbosacral

